# Fatty acid-mediated induction of CYP2E1 activity in HepaRG cells is not systematically associated with exacerbated acetaminophen cytotoxicity

**DOI:** 10.1101/2024.11.01.621521

**Authors:** Clémence Penhoat, Julie Massart, Gregory Pinon, Daniel Catheline, Vincent Rioux, Bernard Fromenty, Karima Begriche

## Abstract

Hepatic cytochrome P450 2E1 (CYP2E1) is thought to contribute to the pathophysiology of metabolic dysfunction-associated steatotic liver disease (MASLD) formerly known as non-alcoholic fatty liver disease (NAFLD). Indeed, increased activity of CYP2E1 in obese subjects with fatty liver may contribute to higher oxidative stress, mitochondrial dysfunction and the progression to steatohepatitis. Besides, higher CYP2E1 activity in obesity and MASLD is deemed to increase the risk of acetaminophen (APAP)-induced hepatotoxicity because this pain reliever is metabolized by CYP2E1 to N-acetyl-p-benzoquinone imine (NAPQI), a highly toxic metabolite. Although the mechanism of MASLD-associated CYP2E1 induction is still unclear, evidence suggests that hepatic CYP2E1 activity is regulated by fatty acids (FAs). In this study, we investigated the effect of 9 FAs differing by their carbon chain length and their degree of unsaturation on CYP2E1 activity in differentiated HepaRG cells. One-week incubation with palmitic acid (PA), stearic acid (SA) and linoleic acid (LA) induced CYP2E1 activity but only LA exposure induced triglyceride accumulation. APAP hepatotoxicity was then assessed in HepaRG cells cultured with or without PA, SA or LA. Acute APAP cytotoxicity was exacerbated in presence of PA or SA and this was accompanied by more severe mitochondrial dysfunction. These effects were not observed when cells were incubated with LA. Hence, FA-mediated increased CYP2E1 activity in HepaRG cells does not necessarily require steatosis. In addition, FA-mediated CYP2E1 induction does not systematically lead to higher APAP-induced cytotoxicity. By favoring triglyceride accumulation, LA might curb APAP-induced mitochondrial dysfunction and cell death.

## INTRODUCTION

Initially described as a fat accumulation in the liver without excessive alcohol consumption, metabolic dysfunction-associated steatotic liver disease (MASLD) formerly known as non-alcoholic fatty liver disease (NAFLD) (Rinella et al., 2023) refers to a large spectrum of liver diseases ranging from metabolic dysfunction-associated steatotic liver (MASL) or simple steatosis to more severe hepatic diseases such as metabolic dysfunction-associated steatohepatitis (MASH), fibrosis, cirrhosis and hepatocellular carcinoma (Lindenmeyer et al., 2018). MASLD concerns about 25% of the worldwide general adult population but this prevalence is significantly higher among patients with obesity or diabetes (Fazel et al., 2016; Younossi., 2019). Most importantly, these two metabolic disturbances were reported to be closely linked to MASH, a severe form of MASLD (Wanless et al., 1990; Leung et al., 2017; Polyzos et al., 2019; Sheka et al., 2020; Stefan and Cusi., 2022). MASH is the inflammatory subtype of MASLD characterized by hepatocyte ballooning and necrosis. MASH can evolve to fibrosis (Brown and Kleiner, 2016) and cirrhosis (Suzuki and Diehl, 2017). These advanced forms of MASLD have been often associated with a greater risk of liver-related morbidity and mortality (Dulai et al., 2017; Taylor et al., 2020; Loomba et al., 2021). Because of these negative clinical outcomes, the understanding of the mechanisms involved in the aggravation of simple fatty liver to MASH appears thus crucial. Cytochrome P450 2E1 (CYP2E1) has been hypothesized as one of the key players in MASH disease since increased hepatic expression and/or activity of this enzyme was reported in different animal models of obesity (Raucy et al., 1991; Begriche et al., 2008; Aubert et al., 2011), steatohepatitis (Weltman et al., 1996) as well as in MASLD /MASH patients (Weltman et al., 1998; Niemelä et al., 2000; Chalasani et al., 2003). The critical role of CYP2E1 in fatty liver aggravation to MASH was subsequently pinpointed in *Cyp2E1* null mice since CYP2E1 deletion provided a protection against high fat diet-induced steatohepatitis (Abdelmegeed et al., 2012). The clinical relevance of CYP2E1 induction on MASLD progression was strongly suggested by the positive correlation between CYP2E1 activity and the degree of steatosis in patients (Emery et al., 2003; Chtioui et al., 2007). Moreover, hepatic CYP2E1 activity might even be higher in obese MASH patients compared to obese individuals with simple steatosis (Orellana et al., 2006). Since CYP2E1 is able to generate significant amounts of reactive oxygen species (ROS) and deleterious endogenous lipid derivates such as dicarboxylic acids (Robertson et al., 2001; Caro et al., 2004; Leung and Nieto., 2013), this cytochrome P450 could then promote the progression of MASLD by triggering an oxidative stress and various related cellular damages such as lipid peroxidation and mitochondrial dysfunction (Begriche et al., 2013; Abdelmegeed et al., 2017; Massart et al., 2022). In obesity-associated MASLD, high circulating levels of fatty acids (FAs), hyperleptinemia and insulin resistance have been reported as possible metabolic inducers of hepatic CYP2E1 (Aubert et al., 2011; Massart et al., 2022). Various experimental studies using high fat-fed rodent models or hepatic cells emphasized the amount of fat intake (Osabe et al., 2008) and most importantly the role of specific FAs in the induction of hepatic CYP2E1 (Yoo et al., 1991; Aubert et al., 2011; Jian et al., 2017; Wang et al., 2020; Raucy et al., 2004; Sung et al., 2004; Michaut et al., 2016; Le Guillou et al., 2018; Bucher et al., 2018; Liu et al., 2019). For instance, *in vitro* investigations carried out on human primary hepatocytes or on differentiated human hepatoma HepaRG cells revealed an enhanced CYP2E1 expression or activity after incubation with palmitic or stearic FAs (Raucy et al., 2004; Michaut et al., 2016; Le Guillou et al., 2018; Bucher et al., 2018). More recently, we showed in HepaRG cells a significant increase in gene expression and activity of CYP2E1 after a 2-week treatment with a mixture of stearate and oleate (Le Guillou et al., 2018; Bucher et al., 2018). Interestingly, this CYP2E1 induction was associated with a moderate overproduction of ROS and mild mitochondrial dysfunction (Bucher et al., 2018).

Besides its potential role in MASLD progression, CYP2E1 plays also an important role in the biotransformation of numerous exogenous compounds such as alcohol, drugs, environmental toxicants and industrial chemicals (Chen et al., 2019; Massart et al., 2022). Regarding some drugs such as the painkiller acetaminophen (APAP, also referred to as paracetamol), CYP2E1 has long been known to be pivotal in the metabolism and liver toxicity of this drug, especially when provided at high doses. In this context, CYP2E1-mediated biotransformation of APAP generates excessive amounts of N-acetyl-*p-*benzoquinone imine (NAPQI) leading to severe liver damage. Indeed, NAPQI is a highly reactive metabolite inducing oxidative stress, mitochondrial dysfunction, ATP depletion and massive necrotic cell death (Ramachandran and Jaescke., 2019; McGill and Hinson., 2020). Although there are some discrepancies between studies in the higher susceptibility to acute APAP hepatotoxicity during metabolic diseases (Ito et al., 2006; Kim et al., 2017; Rutherford et al., 2006; Radosevich et al., 2016), some studies have reported that obesity and MASLD could increase the risk and/or the severity of APAP-induced liver injury in humans (Nguyen et al., 2008; Myers and Shaheen 2009; Chomchai and Chomchai., 2018) and in rodents (Corcoran et al., 1987; Aubert et al., 2012; Kučera et al., 2012; Piccinin et al., 2019; Shi et al., 2019). Acute toxicity of APAP was also investigated in steatotic hepatocytes indicating that exposure to some FAs could exacerbate APAP hepatotoxicity (Michaut et al., 2016; Yang et al., 2020). Among the various factors contributing to the increased susceptibility to acute APAP hepatotoxicity in the context of obesity and MASLD, CYP2E1 induction may be involved in part in this greater toxicity (Aubert et al., 2012; Michaut et al., 2016).

To better understand the role of hepatic CYP2E1 in MASLD pathophysiology and its implication in the susceptibility to acute APAP hepatotoxicity, our present work aims at identify FAs that could induce CYP2E1 in differentiated HepaRG cells, a suitable cell model to investigate MASLD and APAP toxicity (Michaut et al., 2016; McGill et al., 2011). For this purpose, we performed a screening on a set of 9 FAs differing from each other by their carbon chain length and their degree of unsaturation. After a 1-week incubation period with the different FAs, CYP2E1 activity was measured. The 3 FAs able to induce CYP2E1 were then investigated further to determine whether hepatic CYP2E1 induction was associated with triglyceride accumulation. Finally, we determined if the acute hepatotoxicity of APAP could be exacerbated in HepaRG cells incubated with these 3 different FAs. In aggregate, our results indicate that FA-mediated CYP2E1 activity induction favors higher APAP-induced cytotoxicity in HepaRG cells with some but not all FAs, which might be due to their respective ability to promote mitochondrial dysfunction and/or induce cytoprotective mechanisms.

## MATERIALS AND METHODS

### Chemicals and reagents

Acetaminophen (APAP), chlorzoxazone (CZX), cycloheximide (CHX), dimethylsulfoxide (DMSO), L-carnitine, lauric acid (LAU), palmitic acid (PA), stearic acid (SA), oleic acid (OA), linoleic acid (LA), α-linolenic acid (ALA), arachidonic acid (ARA), eicosapentaenoic acid (EPA), docosahexaenoic acid (DHA), fatty acid-free bovine serum albumin (BSA), insulin, protease and phosphatase inhibitors were all purchased from Sigma-Aldrich (Saint-Quentin Fallavier, France). William’s E medium, Dulbecco’s phosphate buffered saline (DPBS), Hanks Balanced Salt Solution (HBSS), glutamine, penicillin, streptomycin, MitoSOX Red probe, dichlorodihydrofluorescein diacetate (H_2_DCFDA) probe, Nile Red, Hoechst 33342, High Capacity cDNA Reverse Transcription Kit (Applied Biosystems), Power SYBR Green PCR Master Mix (Applied Biosystems) and the Pierce bicinchoninic acid (BCA) protein assay kit were all obtained from ThermoFisher Scientific (Courtaboeuf, France). Fetal bovine serum (FBS) was supplied by Lonza (Levallois-Perret, France) and Eurobio Scientific (Les-Ulis, France). Hydrocortisone hemisuccinate was obtained from SERB laboratories (Paris, France). Radiolabeled [U-^14^C] palmitic and [2-^14^C] acetic acids were purchased from PerkinElmer (Villebon-Sur-Yvette, France). RIPA buffer 10X (9806-Cell Signaling) was purchased from Ozyme (Saint-Cyr-L’Ecole, France). Cytochrome P450 2E1 (CYP2E1) antibody (PA26-Oxford Biomedical) was purchased from Euromedex (Souffelweyersheim, France) and Heat-shock cognate 70 (HSC 70) antibody (sc-7298-Santa Cruz Biotechnology) was supplied by CliniSciences (Nanterre, France).

### Cell culture and treatments

The HepaRG cell line was cultured as previously described (Gripon et al., 2002). Briefly, HepaRG progenitors were seeded at a low density of 2.6 × 10^4^ cells/cm^2^ and incubated for two weeks in a growing medium containing William’s E medium supplemented with 10 % FBS, 100 units/mL penicillin, 100 μg/mL streptomycin, 5 μg/mL insulin, and 50 μM hydrocortisone hemisuccinate. Cells were grown at 37°C, 5 % CO_2_ in a humidified incubator and media was refreshed three times a week. When committed to differentiation, HepaRG cells were shifted to a differentiation medium for two additional weeks. The differentiation medium is composed of the growing medium supplemented with 1.75 % DMSO. Once differentiated, HepaRG cells were subsequently treated with different FAs and/or APAP in the differentiation medium slightly modified for FBS (5 %) and DMSO (1 %). For fatty acid treatments, differentiated HepaRG cells were incubated for one week with 150 µM of LAU (C12:0); PA (C16:0), SA (C18:0), OA (C18:1), LA (C18:2), ALA (C18:3). For the long chain FAs namely ARA (C20:4), EPA (C20:5) and DHA (C22:6), the used concentration was 25 µM. For the APAP treatment, differentiated HepaRG cells were treated with this drug at 10 mM or 20 mM during the last 24 h or 48 h.

Of note, all FAs and APAP were dissolved in DMSO and final DMSO concentration in culture media was always set up at 1 % regardless of the treatment. HepaRG cells were used between passage 12 to 16. Unless specified, all investigations were performed 24 h after the last treatment.

### CYP2E1 activity

CYP2E1 activity was measured by determining the production of chlorzoxazone-O-glucuronide (CZX-O-Glc) from chlorzoxazone (CZX) as previously described (Quesnot et al., 2018). Prior to CZX incubation, HepaRG cells were rinsed twice with pre-warmed DPBS and then incubated for 6 h at 37°C in phenol red-free William’s E medium containing 300 µM CZX. At the end of the incubation, aliquots of culture media were harvested and CZX-O-Glc was quantified using a high-performance liquid chromatography (HPLC Agilent Series 1260 Infinity II, Santa Clara, CA). The sample injection volume was 45 µL and the separation was performed with a C18 column (150 mm × 3 mm Accupore PFP, ThermoFisher Scientific). The mobile phase consisted of a mixture of two solvents: solvent A (0.1% of acetic acid + 0.25% of triethylamine hydrochloride) and solvent B (100% of acetonitrile), pumped at a rate of 0.5 mL/min. The gradient program was as follows: 0-1 min: 98% of solvent A and 2% of solvent B; 20-22 min: 10% of solvent A and 90% of solvent B; 27-30 min: 98% of solvent A and 2% of solvent B. The eluates were monitored by an ultraviolet detector at λ= 284 nm. The peak areas of the CZX-O-Glc were automatically integrated using the Agilent 1100 software and quantification was performed from a calibration curve generated with known amounts of CZX-O-Glc. CYP2E1 activity was expressed as pmol CZX-O-Glc/mg of proteins/hour.

### RNA extraction and gene expression

Total RNAs were extracted from about 6 ×10^6^ HepaRG cells using a Nucleospin RNA isolation Kit (Macherey-Nagel, Hoert, France) following the manufacturer’s instructions. After RNA quantification with the NanoDrop One (ThermoFisher Scientific, Courtaboeuf, France), cDNAs were synthesized from 500 ng of total RNA using the High Capacity cDNA Reverse Transcription Kit (Applied Biosystems). cDNAs were then amplified with specific primers using the Power SYBR Green PCR Master Mix (Applied Biosystems) and a 384-well QuantStudio™ 7 Flex Real-Time PCR System (Thermo Fisher Scientific). The primer sequences used for quantitative PCR are detailed in Table 1. Expression of the human *CYCLOPHILIN B gene* was used as a reference, and the 2 ^ΔΔT^ method was used to express the relative expression of each selected gene. Non-treated cells were used as a control group for gene expression which was arbitrarily set at 1. To determine if the expression of the *CYP2E1* gene is under the direct control of the different FAs, cloning of *CYP2E1* promoter and transcriptional assay were conducted as described in the Supplementary data.

### Protein extraction and Western Blot analysis

Prior to cell lysis, media was removed and HepaRG cells were rinsed twice in ice-cold DPBS buffer. Then, cells were lysed in ice-cold 1X RIPA buffer supplemented with protease and phosphatase inhibitors. Protein concentration was determined using the Pierce BCA protein assay kit as recommended by the manufacturer’s instructions. Equal amounts of protein (25 µg) were diluted with Laemmli buffer and heated at 56°C for 20 minutes before being separated by SDS-PAGE using NuPAGE 4-12 % Bis-tris gels (ThermoFisher Scientific, Courtaboeuf, France) and then transferred to 0.2 µM nitrocellulose membrane (Bio-Rad, Les Ulis, France). Ponceau staining was used to confirm equal loading and transfer. Membranes were then blocked in 7.5 % skimmed dry milk in TBST (10 mM Tris-HCl, 100 mM NaCl, 0.03 % Tween 20) for 1 hour at room temperature and incubated overnight with CYP2E1 antibody (dilution 1:2000) or HSC70 (dilution 1:500). After three successive wash steps in TBST, the membrane was incubated with an appropriate HRP-conjugated secondary antibody and protein bands were revealed by enhanced chemiluminescence (Pierce SuperSignal West Dura Extended Duration Substrate, ThermoFisher Scientific) and visualized using Fusion FX imaging system (Vilber Lourmat, Marne La Vallée, France). Protein content was quantified by densitometry with ImageJ software (National Institutes of Health, Bethesda, MD). HSC70 protein content was used for normalization.

### Effect of cycloheximide on CYP2E1 protein expression

To assess mechanisms involved in CYP2E1 protein expression regulation, HepaRG cells were treated or not with 150 µM of PA, SA or LA for 1 week. To follow CYP2E1 protein expression over time, cells were subsequently incubated with cycloheximide (50 µM), a protein synthesis inhibitor for 48 h, 72 h and 96 h. CYP2E1 protein content was determined by Western blot for each time point. CYP2E1 protein expression was normalized to Ponceau staining intensity around CYP2E1 area. To display variation in CYP2E1 protein expression results were expressed as Δ_t(96)-t (0)_ CYP2E1/Ponceau.

### Nile Red staining, intracellular triglyceride content and apolipoprotein B secretion

Nile Red staining was used to visualize neutral lipids accumulation within HepaRG cells after a one-week treatment with the different FAs. As previously described (Bucher et al., 2018), cells were first rinsed with pre-warmed DPBS and fixed with 4 % paraformaldehyde for 20 minutes at room temperature. Cells were then washed three times with DPBS and incubated with 0.1 µg/mL Nile Red in DPBS for 30 minutes at room temperature. After a washing step, cells were incubated with 10 µg/mL Hoechst 33342 in DPBS for 15 minutes at room temperature to label nuclei. Intracellular accumulation of neutral lipids was assessed using fluorescent microscope (Axio Vert.A1 and Colibri.2 controller, Zeiss) with the appropriate excitation/emission wavelengths respectively 531/461nm for Nile Red and 531/593nm for Hoechst 33342. To determine triglyceride content within HepaRG cells, the triglyceride quantification kit from BioVision was used (Waltham, MA). Briefly, HepaRG cells were homogenized in 5 % NP-40 detergent to allow triglyceride extraction. Then, to completely solubilize triglycerides, samples were heated at 100°C for 5 minutes and centrifuged. Solubilized triglycerides were collected and then hydrolyzed to FAs and glycerol by adding lipase. Lipase was also added to each triglyceride standard aliquot. According to the manufacturer’s instructions, free glycerol was then oxidized in a specific mixture containing a probe. After 30 minutes of incubation at room temperature, the newly formed oxidized product reacts with the probe to generate a colored derivative. This derivative was then measured at λ= 570 nm using a FLUOstar omega microplate reader (BMG Labtech, Ortenberg, Germany). Of note, the triglyceride content of each sample was determined by comparison to a standard curve generated with known amounts of triglycerides. The methods used to determine the fatty acid composition of lipids and triglycerides are provided in the Supplementary data.

To assess ApoB secretion, culture media was harvested at the end of the 7 days of treatment with the different FAs. The Human Apolipoprotein B Elisa kit from Mabtech (Nacka Strand, Sweden) was used for ApoB quantification and as previously described (Allard et al., 2021).

### De novo lipogenesis and mitochondrial β-oxidation

*De novo* lipogenesis was assessed in adherent HepaRG cells by measuring newly synthetized lipids from [2-^14^C] acetic and as previously described (Allard et al., 2021). Mitochondrial fatty acid oxidation (mtFAO) was determined by measuring the acid-soluble radiolabeled metabolites resulting from mitochondrial oxidation of [U-^14^C] palmitate. Briefly, adherent HepaRG cells in 96-well plate were incubated for 3 hours at 37°C in a red-free William’s E medium supplemented with 1 % fatty acid-free BSA, 1 mM L-carnitine, [U-^14^C] palmitic acid (185 Bq/well), 100 µM cold PA and 1 % DMSO. At the end of the incubation, 100 µL of 6 % perchloric acid was directly added to each well in order to precipitate non-oxidized fatty acids. Then, cells were centrifuged at 2000*g* for 10 minutes at room temperature. After centrifugation, 130 µL of supernatant was collected and transferred in 4 mL of scintillation liquid. [U-^14^C] labeled acid-soluble β-oxidation products were measured in a Tri-Carb 4910TR liquid scintillation counter (PerkinElmer, Villebon-Sur-Yvette, France). Results were normalized to total protein content and expressed as count per minute (CPM)/ µg of proteins.

### Assessment of intracellular ATP levels

The intracellular content of ATP was measured using the CellTiter Glo Luminescent cell viability assay kit from Promega (Charbonnières-Les-Bains, France). Of note, ATP quantification reflects cell viability since ATP is produced from metabolically active cells. To assess cell viability in response to FAs and APAP treatments, differentiated HepaRG cells were first incubated for 1 week with 150 µM of PA, SA or LA. The last 24 or 48 hours, some cells were co-treated or not with 10 or 20 mM APAP. Before performing the ATP assay, HepaRG cells were first rinsed twice with pre-warmed DPBS and kept for 30 minutes at room temperature in phenol red-free William’s E medium to equilibrate plate temperature and avoid any inconsistent results due to temperature differences. Cells were subsequently incubated with the CellTiter-Glo reagent mix for 10 minutes at room temperature. At the end of the incubation, HepaRG cells were transferred in a white opaque 96-well microplate and the emitted luminescent signal was quantified using a FLUOstar Omega microplate reader (BMG Labtech, Ortenberg, Germany). Results were expressed in comparison to control cells.

### Assessment of reactive oxygen species (ROS) production

To measure ROS production, H_2_DCFDA and MitoSox Red fluorescent probes were used for the detection of intracellular hydrogen peroxide and mitochondrial superoxide anion, respectively. Prior to the assay, HepaRG cells were treated for one week with 150 µM of PA, SA or LA. During the last 24 hours, some cells were co-incubated or not with 20 mM APAP. The day of the assay, HepaRG cells were treated one more time for 15 or 90 minutes with APAP alone or in combination with the different FAs. At the end of the kinetic, medium was quickly removed and HepaRG cells were rinsed once with pre-warmed HBSS. Then, cells were incubated for 30 minutes at 37°C in HBSS supplemented with 2 µM of H_2_DCFDA or 5 µM of MitoSox Red probes. At the end of the incubation, living cells were gently rinsed and fluorescence was measured using a FLUOstar Omega microplate reader (BMG Labtech, Ortenberg, Germany) with excitation/emission wavelengths of 485/520 nm for H_2_DCFDA and 520/590 nm for MitoSOX Red. Fluorescence intensity was normalized to total protein content and results were expressed in comparison to control cells.

### Assessment of mitochondrial respiration

Mitochondrial respiration was assessed by measuring the oxygen consumption rate (OCR) using the Seahorse XFe24 flux analyzer and XF Cell Mito Stress test kit (Agilent, Santa Clara, CA) and according to the manufacturer’s instructions. Briefly, HepaRG cells were seeded and differentiated in the XFe24 cell culture plate (Agilent, Santa Clara, CA). Once differentiated, cells were treated or not for 1 week with 150 µM of PA, SA or LA. Some cells were co-treated with 20 mM APAP during the last 24 h. On the day of the assay, the medium was removed and replaced with a bicarbonate-free low-buffered medium (Seahorse Bioscience) containing 10 mM glucose, 2 mM glutamine, 1 mM sodium pyruvate (pH 7.4). After a 45 minutes incubation at 37°C in the absence of CO_2_, cells were then placed into the XFe24 flux analyzer, and the OCR was evaluated by sequential injection of 2 μM oligomycin, 2 μM carbonyl cyanide-4-trifluoromethoxy phenylhydrazone (FCCP), and 0.5 μM rotenone/antimycin A. Then, the OCR provided by the Seahorse Mito Stress Test profile and corresponding to basal respiration, maximal respiration (induced by the OXPHOS uncoupler FCCP) and respiration linked to ATP production and spare respiratory capacity were normalized in each well to the number of cells estimated by the fluorescence intensity of the Hoechst 33342 dye (10 μg/ml) using the FLUOstar Omega microplate reader from BMG Labtech (Ortenberg, Germany) with excitation/emission wavelengths of 355/460 nm.

### Statistical analysis

Statistical analyses were performed using GraphPad Prism 8.0.2 software. All results are expressed as mean ± SEM (standard error of mean). Non-parametric test was used. The significance was evaluated by Mann Whitney test. Comparisons were considered statistically significant at *p* ≤ 0.05.

## RESULTS

### Identification of FAs inducing hepatic CYP2E1 and associated mechanisms

In order to identify FAs that could induce CYP2E1 in differentiated HepaRG cells, we performed a screening with a set of 9 FAs differing from each other by their carbon chain length and degree of unsaturation. The selected FAs were LAU (C12:0), PA (C16:0), SA (C18:0), OA (C18:1), LA (C18:2), ALA (C18:3), ARA (C20:4), EPA (C20:5) and DHA (C22:6). First, we assessed intracellular ATP content following 1-week treatment for each of these FAs using a wide range of concentrations (i.e 10, 25, 50, 100, 150, 200, 300 and 400 µM) to determine their potential cytotoxicity. HepaRG cells incubated for 1 week with PA, SA or ALA exhibited significant loss of ATP when FA concentration was equal to or greater than 200 µM whereas no effect was observed for LAU and OA (data not shown). For ARA, EPA and DHA, treatment with concentrations greater or equal to 50 µM induced significant cell toxicity (data not shown). We thus chose for our study to use 150 µM for medium and long-chain FAs (LAU, PA, SA, OA, LA and ALA) and 25 µM for long-chain polyunsaturated FAs (ARA, EPA and DHA). Of note, these two concentrations are within the range of physiological circulating levels of most non-esterified FAs with a carbon chain length varying from C12 to C22 (Kangani et al., 2008; Puri et al., 2009).

We next determined CYP2E1 activity in differentiated HepaRG cells treated for 1 week with each of the 9 aforementioned FAs. In these experimental conditions, PA, SA and LA were able to significantly increase the activity of CYP2E1 by 59 %, 44 % and 28 % respectively **(Figure 1)**. Conversely, CYP2E1 activity dropped in HepaRG cells incubated with ALA **(Figure 1)**. CYP2E1 activity remained unchanged following 1-week treatment with LAU, OA, ARA EPA or DHA **(Figure 1)**. Since MASLD is most commonly associated with an induction of CYP2E1 (Aubert et al., 2011; Begriche et al., 2023), FAs that increased CYP2E1 activity (i.e. PA, SA and LA) were further investigated. Induction of CYP2E1 could result from enhanced gene expression, increased translation or protein stability. We thus measured CYP2E1 mRNA and protein levels after 1-week treatment with 150 µM PA, SA or LA. However, these FAs did not increase *CYP2E1* gene expression in HepaRG cells compared to untreated control cells **(Figure 2 A)**. Additional molecular investigations based on a luciferase assay reporting the activity of CYP2E1 promoter were performed in HepG2 cells. Cells transfected with the entire human promoter were then exposed to PA, SA or LA for 2, 4, 6, 8, 12 or 24 h. Luciferase reporter assays showed that none of the 3 FAs caused *CYP2E1* promoter transactivation **(Supplemental Figure 1)**. Thus, these results indicated that neither PA, SA nor LA had a direct effect on *CYP2E1* gene expression **(Supplemental Figure 1)**. Similar to our mRNA and luciferase assay results, CYP2E1 protein levels were not significantly increased, despite a trend after a 1-week incubation with PA, SA or LA (Figure 2 B). In order to investigate if PA, SA or LA could induce CYP2E1 activity through a post-transcriptional mechanism, we next treated HepaRG cells with PA, SA or LA for 1 week in presence of cycloheximide for the last 48 h, 72 h or 96 h of the treatment. As expected, CYP2E1 protein expression in HepaRG cells progressively declined with protein synthesis inhibition. Interestingly, this decrease tended to be attenuated when HepaRG cells were cultured with PA, SA or LA **(Figure 2 C).** After a 96-h co-incubation with cycloheximide, CYP2E1 protein expression decreased by half in untreated control cells whereas this reduction was less marked in HepaRG cells treated with PA, SA or LA, although not statistically significant (Figure 2 C). Hence, these results suggest that PA, SA or LA-mediated CYP2E1 induction might be due to less degradation and/or stabilization of CYP2E1 at the protein level, contributing to higher activity.

**Figure 1:**
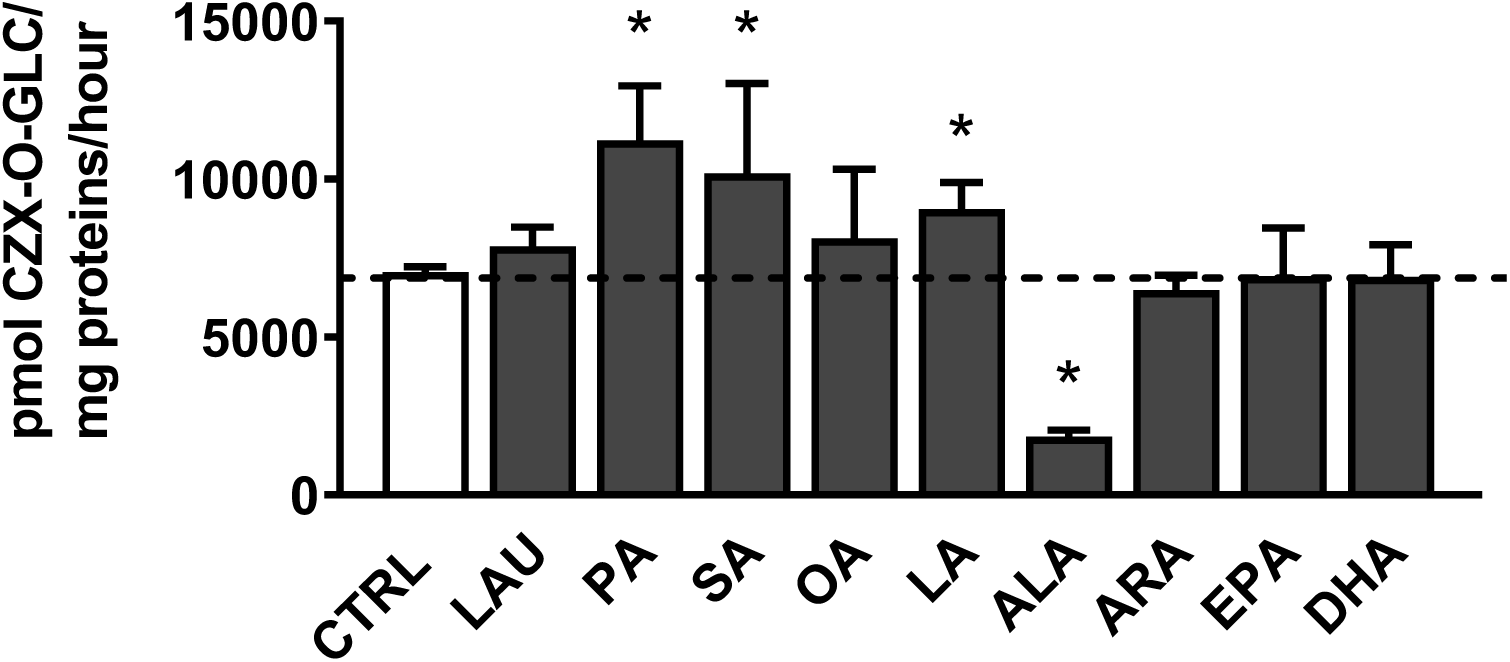
CYP2E1 activity in HepaRG cells treated for 1 week with various fatty acids. Differentiated HepaRG cells were incubated for 1 week with 150 µM of lauric (LAU), palmitic (PA), stearic (SA), oleic (OA), linoleic (LA) or α-linolenic (ALA) acids or with 25 µM of arachidonic (AA), eicosapentaenoic (EPA) and docosahexaenoic (DHA) acids. Results are expressed as mean ± SEM from 4 independent cell cultures. The horizontal dashed line represents the level of vehicle-treated control cells (CTRL). * significantly different from control cells (CTRL) (p≤0.05)

**Figure 2:**
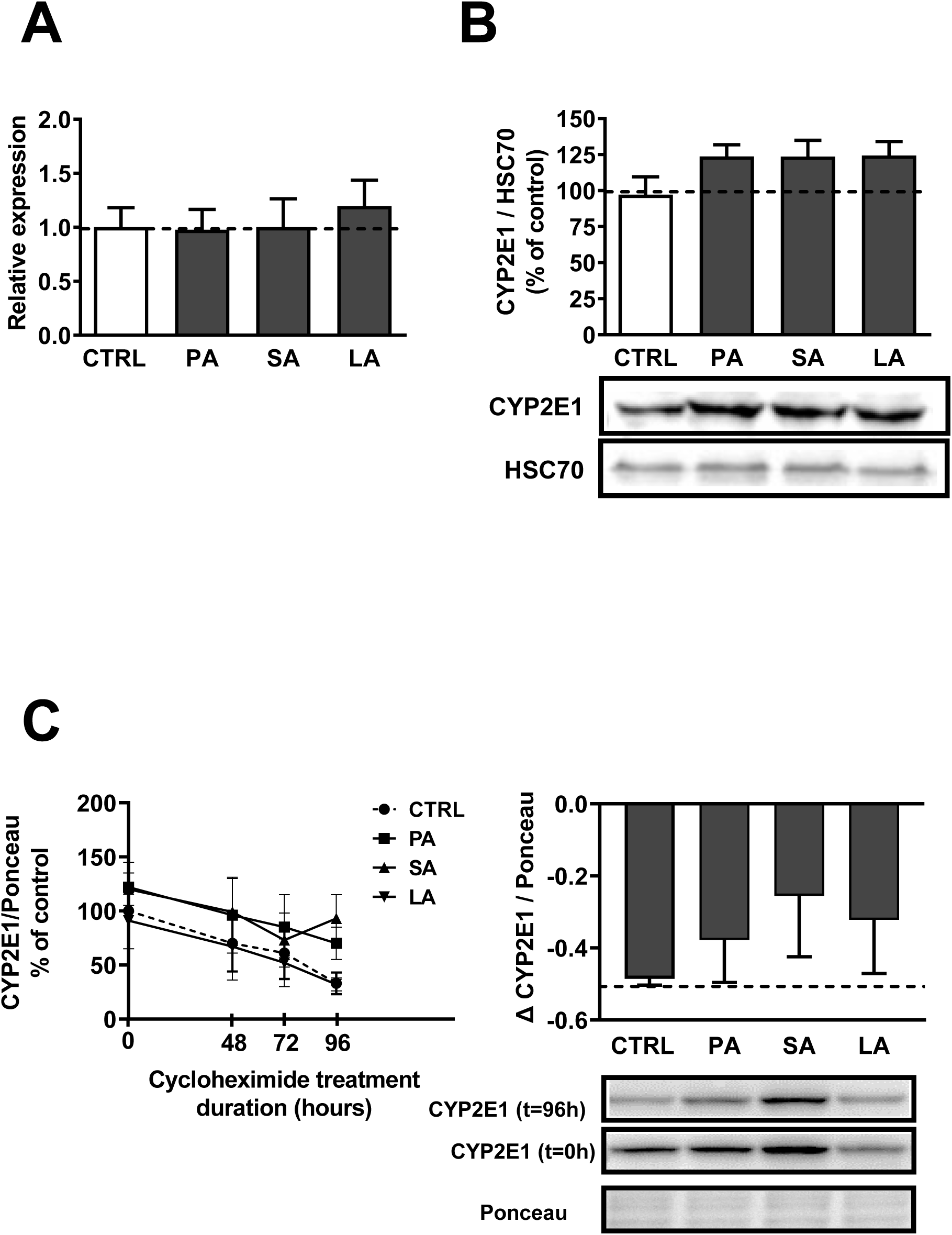
CYP2E1 expression in HepaRG cells treated for 1 week with palmitic, stearic or linoleic acids. Differentiated HepaRG cells were incubated for 1 week with 150 µM of palmitic (PA), stearic (SA) or linoleic (LA) acids. (A) Gene expression of *CYP2E1* determined by RT-qPCR (n=7). (B) Western blot analysis of CYP2E1 (n=4) with a representative picture. (C) Time course of CYP2E1 protein expression in response to cycloheximide treatment (n=5) and relative decrease in expression over the 96 h time course. Results are expressed as mean ± SEM. n indicates the number of independent cell cultures. The dashed line represents the level of control cells (CTRL).

### Assessment of intracellular lipids and investigations on metabolic pathways regulating hepatic lipid homeostasis

Neutral lipids stored within the HepaRG cells were first visualized by fluorescence microscopy using Nile Red staining, a dye used to label fat accumulation in the cytosol (McMillian et al., 2001; Bucher et al., 2018). After 1 week of treatment, only HepaRG cells supplemented with LA exhibited an increase in the intracellular content of neutral lipids compared to untreated control cells **(Figure 3 A).** HepaRG cells incubated with PA or SA had a weak signal equivalent to the one observed in control cells. We further confirmed these observations by direct quantification of intracellular triglycerides. Indeed, triglyceride content was significantly increased in HepaRG cells cultured with LA while unchanged under PA or SA supplementation **(Figure 3 B).** A targeted lipidomic analysis allowing us to assess the fatty acid composition of the lipid and triglyceride fractions confirmed the greater content of total lipids and triglycerides in HepaRG cells under LA supplementation **(Supplemental Figure 2 A-B).** Moreover, the data indicated higher incorporation of LA into triglycerides in HepaRG cells supplemented for 1 week with LA **(Supplemental Figure 2 B).**

**Figure 3:**
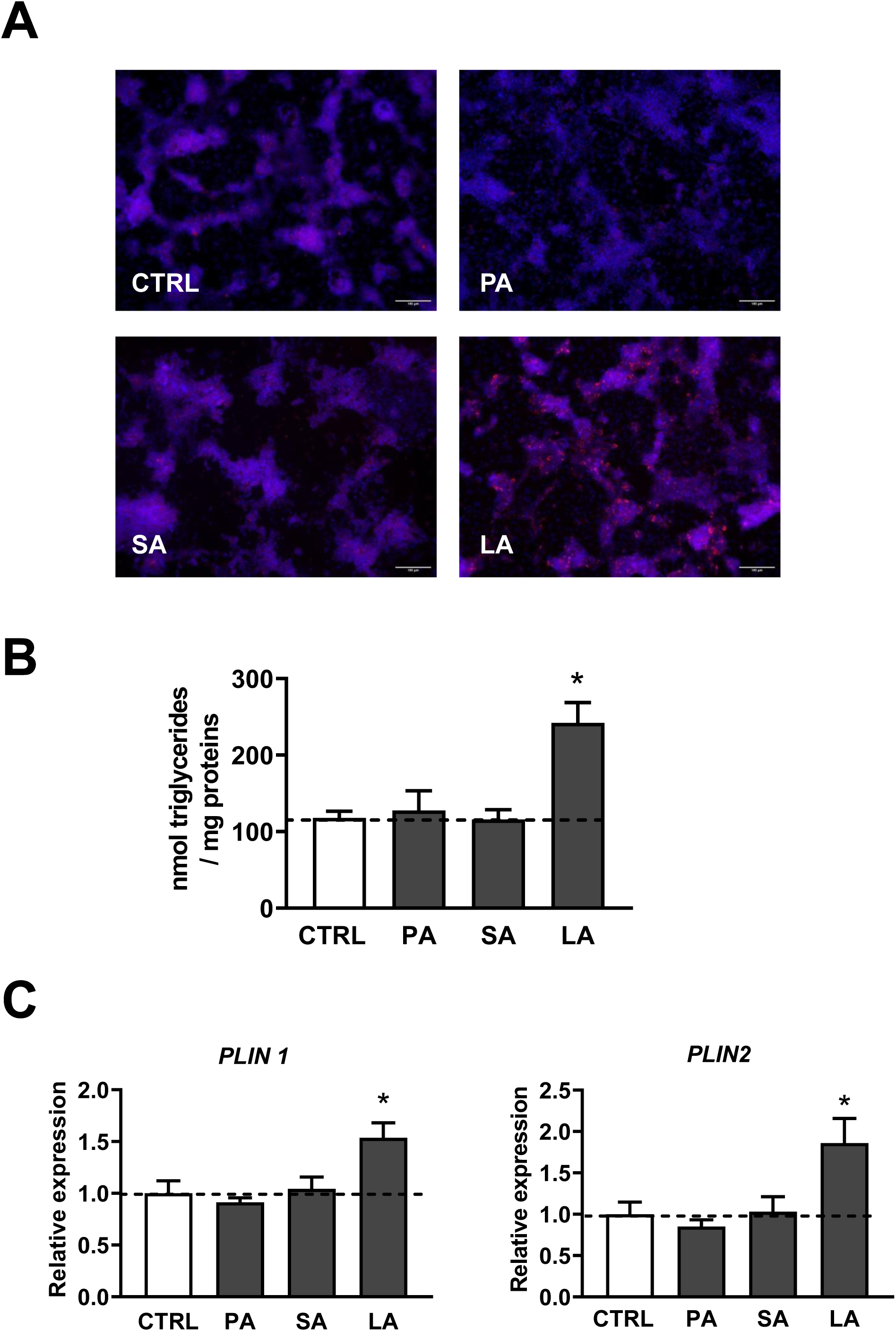
Steatosis in HepaRG cells treated for 1 week with palmitic, stearic or linoleic acids. Differentiated HepaRG cells were incubated for 1 week with 150 µM of palmitic (PA), stearic (SA) or linoleic (LA) acids. (A) Cellular neutral lipids stained with Nile Red and Hoescht. (B) Cellular levels of triglycerides (n=5). (C) Gene expression of perilipin 1 (*PLIN1*) and perilipin 2 (*PLIN 2*) determined by RT-qPCR (n=7). Results are expressed as mean ± SEM. n indicates the number of independent cell cultures. The dashed line represents the level of control cells (CTRL). * significantly different from control cells (CTRL) (p≤0.05).

In order to further characterize lipid accumulation, we investigated the mRNA levels of some lipid droplet-associated genes such as perilipin 1 (PLIN1) and perilipin 2 (PLIN2) as relevant biomarkers of steatosis as previously described (Pawella et al., 2014). Consistent with lipid content, both *PLIN1* and *PLIN2* expression was significantly enhanced after LA incubation, whereas it remained unchanged after PA or SA exposure **(Figure 3 C).** All these results indicate that only LA induces steatosis in our experimental conditions. Thus, FA-induced steatosis in HepaRG cells is not systematically associated with CYP2E1 induction, as such induction was also observed under PA and SA supplementation that did not promote intracellular triglyceride accumulation. Therefore, induction of CYP2E1 in HepaRG cells might rely on FAs themselves and their downstream intracellular effects rather than on the presence of steatosis *per se*.

*De novo* lipogenesis, mtFAO, fatty acid uptake and very low-density lipoprotein (VLDL) secretion are the four major pathways contributing to lipid homeostasis in the liver. A disruption in one or more of these pathways can promote the development of MASLD (Ipsen et al., 2018; Loomba et al., 2021). To determine whether these metabolic pathways were impacted by PA, SA or LA, we first measured gene expression of the fatty acid translocase CD36 (FAT/CD36), a pivotal transporter mediating the uptake of long-chain fatty acids in hepatocytes. Hepatic expression of *FAT/CD36* was increased in a similar manner to the triglyceride content in HepaRG cells **(Figure 4 A)**. Indeed, expression of this transporter was significantly enhanced in HepaRG cells supplemented with LA, which was consistent with the increased intracellular triglyceride content induced by this FA **(Figure 4 A)**. These data are in line with increased expression of FAT/CD36 reported in rodent and human MASLD (Greco et al., 2008; Miquilena-Colina et al., 2011). In contrast, HepaRG cells cultured with PA, and to a non-significant extent with SA, showed a reduced expression of *FAT/CD36* gene expression in comparison to control cells **(Figure 4 A)**.

**Figure 4:**
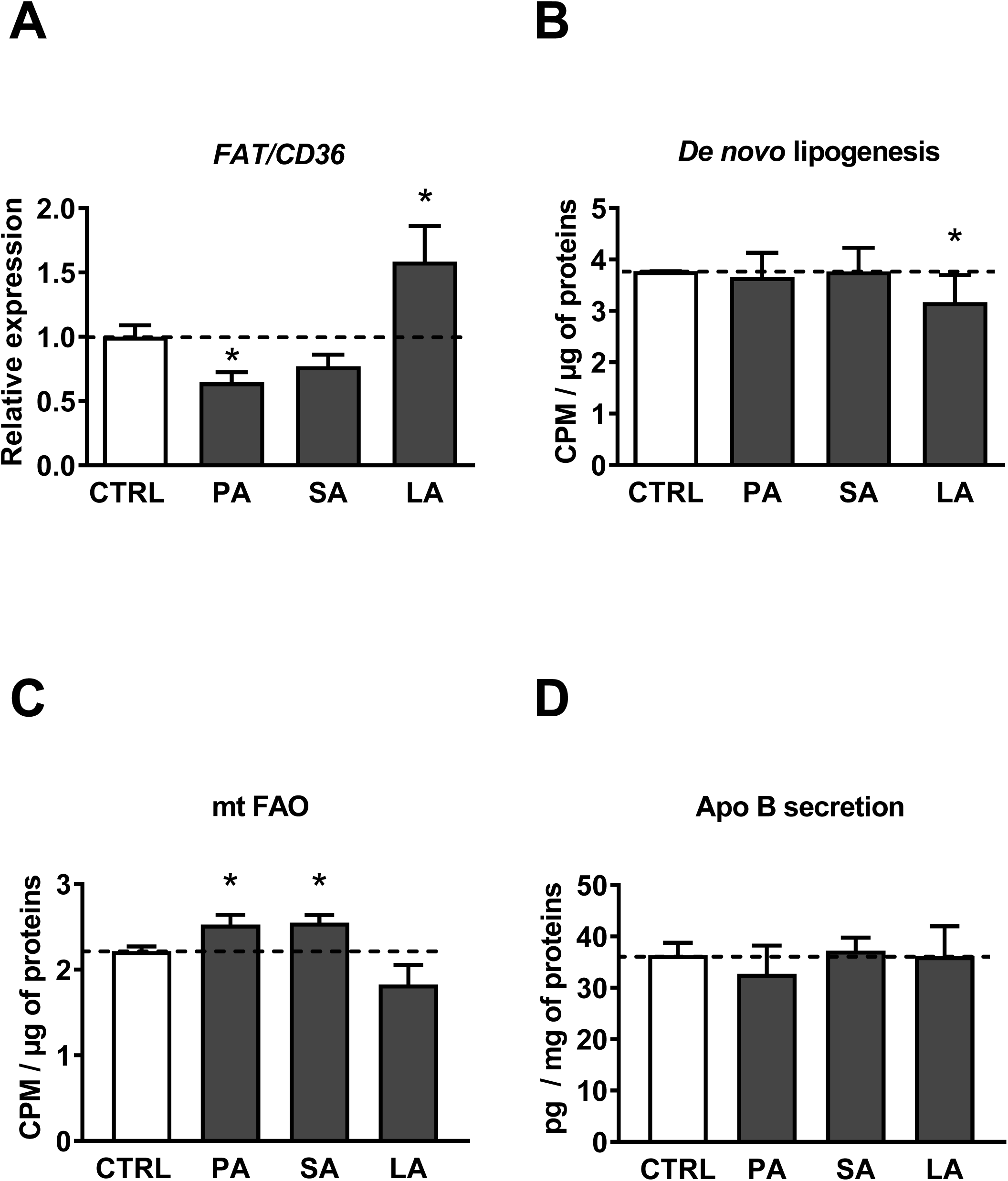
*FAT/CD36* gene expression, *de novo* lipogenesis, mitochondrial fatty acid oxidation and ApoB secretion in HepaRG cells treated for 1 week with palmitic, stearic or linoleic acids. Differentiated HepaRG cells were incubated for 1 week with 150 µM of palmitic (PA), stearic (SA) or linoleic (LA) acids. (A) *FAT/CD36* gene expression (n=7). (B) *De novo* lipogenesis (n=7). (C) Mitochondrial fatty acid oxidation (n=7). (D) ApoB secretion (n=3). Results are expressed as mean ± SEM. n indicates the number of independent cell cultures. The dashed line represents the level of control cells (CTRL). * significantly different from control cells (CTRL) (p≤0.05).

*De novo* lipogenesis was slightly but significantly decreased in HepaRG cells incubated with LA compared to control cells, whereas PA or SA supplementation had no effect **(Figure 4 B)**. Of note, downregulation of lipogenesis is generally observed in response to high fat intake as a compensatory mechanism in order to curb lipid overload (Ren et al., 2012; Allard et al., 2021). In our experimental conditions, the maintenance of a basal rate of lipogenesis despite PA or SA supplementation was probably linked to reduced FA uptake as suggested by decreased *FAT/CD36* expression. We also assessed the expression of some key lipogenic genes, namely sterol regulatory element binding protein 1c (SREBF1, coding for SREBP1c), fatty acid synthase (FASN), acetyl-CoA carboxylase 1 (ACACA, coding for ACC1) and glycerol-3-phosphate acyltransferase mitochondrial isoform (GPAM). A trend for reduced *SREBP1c* expression was observed following FA treatment, which was in line with decreased expression of its target genes *FASN* and *ACC1* **(Supplemental Figure 2 C)**. Interestingly, *GPAM* expression was decreased only following PA and SA exposure **(Supplemental Figure 2 C)**, the 2 FAs that did not induce lipid accumulation.

For mtFAO, a mild but significant increase was measured in HepaRG cells cultured with PA (+14%) or SA (+15%) compared to control cells while LA treatment was without effect **(Figure 4 C)**. Since fatty acid oxidation can be controlled in part by peroxisome proliferator-activated receptor alpha (PPARα), we performed additional investigations to study gene expression of this nuclear receptor and some of its target genes such as carnitine palmitoyl transferase 1 (CPT1) and acyl-CoA oxidase 1 (ACOX1). The expression of all these genes was unchanged regardless of the FA treatment **(Supplemental Figure 3 A)**. These results suggest that increased mitochondrial β-oxidation under PA and SA supplementation might result from non-transcriptional mechanisms, although further investigations would be needed to confirm this hypothesis.

Finally, we investigated VLDL secretion by measuring apolipoprotein B (ApoB) levels in the culture medium as well as the expression of genes encoding ApoB and microsomal triglyceride transfer protein (MTTP), which is a key enzyme for the lipidation of ApoB. However, we did not observe any variation in ApoB secretion **(Figure 4 D)** as well as in *ApoB* and *MTTP* gene expression regardless of the FA treatment **(Supplemental Figure 3 B).**

In aggregate, our data suggest that LA supplementation might induce lipid accumulation in HepaRG cells through increased FA uptake. On the other hand, both PA and SA are not steatogenic in our experimental condition, possibly due to increased mtFAO and reduced *GPAM* expression. Of note, although *de novo* lipogenesis from [2-^14^C] acetic acid was unchanged with PA and SA supplementation lower *GPAM* expression is expected to decrease the generation of triglycerides from glycerol-3-phosphate and neosynthesized FAs.

### APAP induced cytotoxicity, oxidative stress and mitochondrial dysfunction

As mentioned above, some studies argue that hepatic CYP2E1 induction may be involved at least in part in the greater acute hepatotoxicity of APAP in obesity and MASLD (Aubert et al., 2012; Begriche et al., 2023; Michaut et al., 2014). Previously, we reported that SA-dependent CYP2E1 induction promoted higher APAP cytotoxicity in the HepaRG cell line (Michaut et al., 2016). To determine whether PA and LA, two newly identified FAs mediating CYP2E1 induction, also exacerbated APAP cytotoxicity, we incubated HepaRG cells with 150 µM of PA, SA and LA for 1 week and treated with 10 mM or 20 mM APAP the last 24 h or 48 h of the treatment. APAP cytotoxicity was assessed by quantification of intracellular ATP levels. As expected, APAP induced toxicity in HepaRG cells in comparison to control cells not exposed to this drug **(Figure 5)**. Overall, this cytotoxicity increased with APAP concentration and exposure time. Under FAs supplementation conditions, APAP cytotoxicity was more pronounced in HepaRG cells incubated with PA or SA and exposed to 20 mM APAP for 24 h, although the loss of ATP was not statistically different from cells treated only with APAP **(Figure 5 B)**. However, after longer exposure to APAP (48 h), ATP levels in HepaRG cells incubated with PA or SA were significantly decreased compared to HepaRG cells treated with APAP alone **(Figure 5 C and D)**. HepaRG cells supplemented with LA exhibited reduced early cytotoxicity to APAP as decreased ATP levels were not significantly different from control cells after a 24 h exposure to 10 mM or 20 mM APAP **(Figure 5 A and B)**. However, for longer exposure (i.e. 48 h) LA did not affect APAP cytotoxicity as ATP levels in HepaRG cells cultured with LA were similar to HepaRG cells treated with APAP alone **(Figure 5 C and D)**. Collectively, these results indicated that only PA and SA exposures exacerbated APAP cytotoxicity in HepaRG cells whereas LA did not. Thus, FA-mediated CYP2E1 activity induction in HepaRG cells did not systematically lead to higher APAP cytotoxicity. This might suggest the contribution of other factors such as the extent of oxidative stress and mitochondrial dysfunction, which are key events in the initiation of APAP-induced cell death (Ramachandran and Jaescke, 2019; McGill and Hinson, 2020).

**Figure 5:**
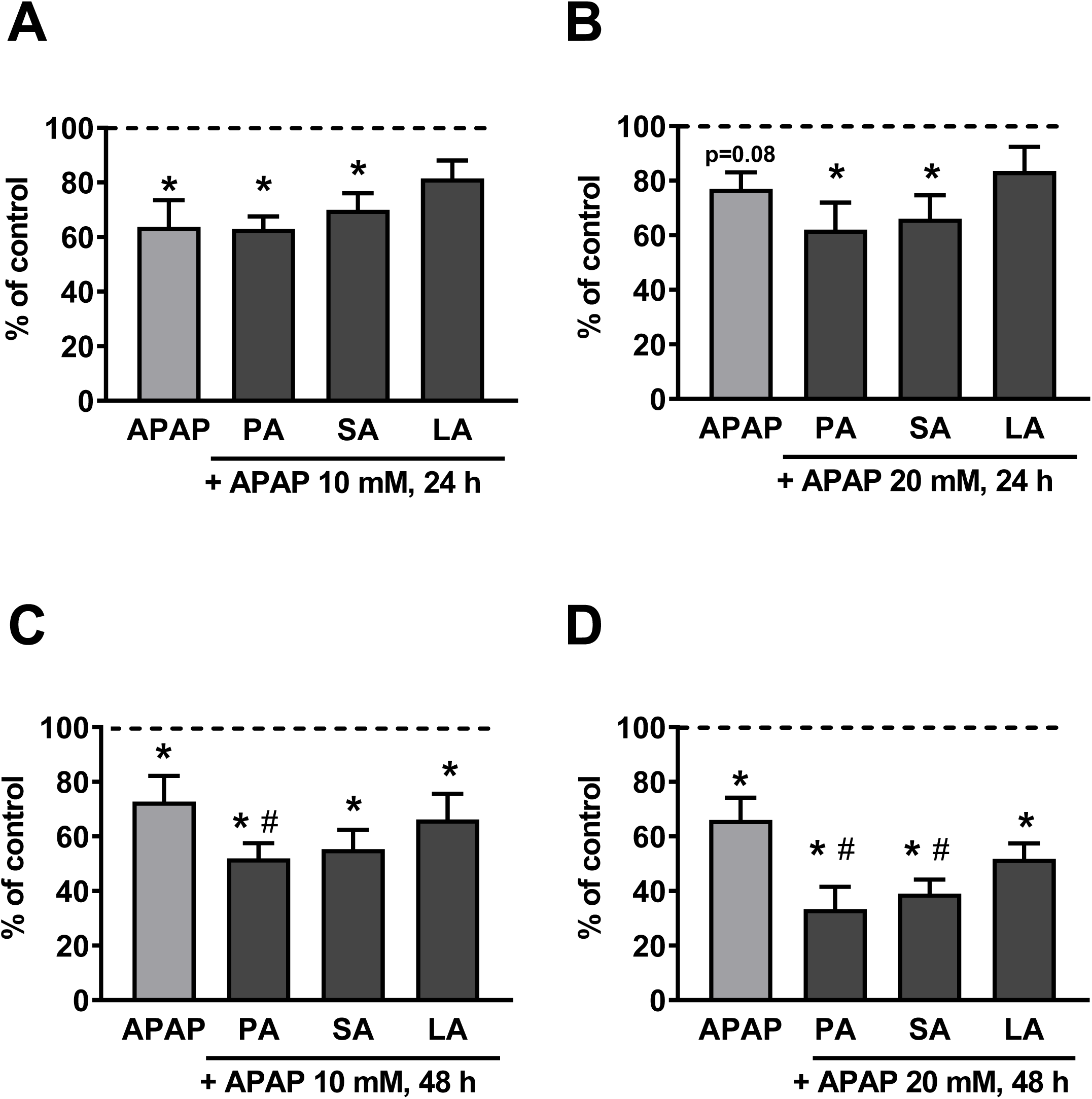
Acetaminophen toxicity in HepaRG cells treated for 1 week with palmitic, stearic or linoleic acids. Differentiated HepaRG cells were incubated for 1 week with 150 µM of palmitic (PA), stearic (SA) or linoleic (LA) acids and exposed to 10 mM or 20 mM acetaminophen (APAP) for the last 24 h or 48 h of the treatment. APAP toxicity was assessed by quantification of intracellular ATP levels. ATP levels after 10 mM (A) or 20 mM (B) APAP treatment for 24 h (n=5). ATP levels after 10 mM (C) or 20 mM (D) APAP treatment for 48 h (n=5). Results are expressed as mean ± SEM. n indicates the number of independent cell cultures. The dashed line represents the level of control cells (CTRL). * significantly different from control cells (CTRL) (p≤0.05). # significantly different from HepaRG cells treated with APAP and without fatty acids (p≤0.05).

Hence, ROS production was first assessed in HepaRG cells as described in the Methods in an attempt to detect early oxidative stress that might explain the exacerbated APAP-induced cytotoxicity observed after 48 h. In absence of APAP, hydrogen peroxide production was enhanced in HepaRG cells cultured with SA and LA but not in cells treated with PA **(Figure 6 A-B)**. However, none of the tested FAs impacted the production of mitochondrial superoxide anion **(Figure 6 C-D)**. After 20 mM APAP, levels of both hydrogen peroxide and superoxide anion increased quickly in all groups of treatment **(Figure 6)**. Yet, LA induced higher hydrogen peroxide levels compared to HepaRG cells treated with APAP alone, while SA induced higher superoxide anion production compared to HepaRG cells treated only with APAP **(Figure 6)**. Collectively, these data suggested that higher APAP-induced cytotoxicity observed with PA and SA **(Figure 5D)** could not be explained by increased oxidative stress since LA (which did not favor APAP cytotoxicity) also enhanced ROS production to a degree similar to or greater than the other two FAs.

**Figure 6:**
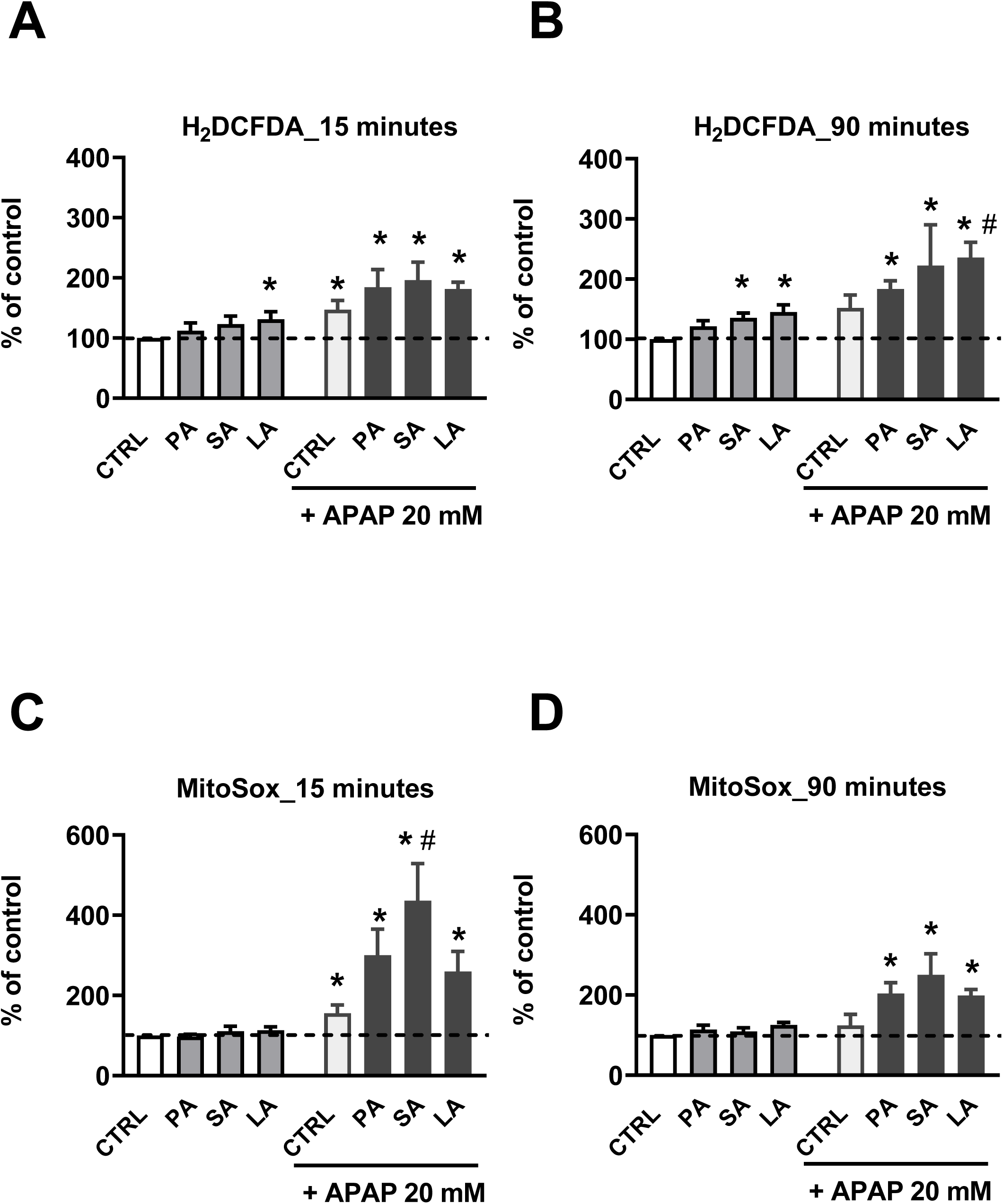
Oxidative stress in HepaRG cells treated for 1 week with palmitic, stearic or linoleic acids and exposed or not to acetaminophen. Differentiated HepaRG cells were incubated for 1 week with 150 µM of palmitic (PA), stearic (SA) or linoleic (LA) acids and exposed or not to 20 mM acetaminophen (APAP) for the last 24 h of the treatment. After this 24 h period, HepaRG cells were treated one more time for 15 or 90 minutes with APAP alone or in combination with the different FAs and ROS were measured 15 or 90 minutes later with H_2_DCFDA and MitoSox probes that detect intracellular hydrogen peroxide and mitochondrial superoxide anion, respectively. Hydrogen peroxide levels at 15 minutes (A) and 90 minutes (B) (n=5). Mitochondrial superoxide anion levels at 15 minutes (C) and 90 minutes (D) (n=5). Results are expressed as mean ± SEM. n indicates the number of independent cell cultures. The dashed line represents the level of control cells (CTRL). * significantly different from control cells (CTRL) (p≤0.05). # significantly different from HepaRG cells treated with APAP and without fatty acids (p≤0.05)

Consequently, mitochondrial respiration was also assessed in the different groups of treatment, as described in the Methods. Without APAP treatment, OCR was not affected by FA exposure, despite a trend for decreased maximal respiration with SA exposure **(Figure 7B)**, indicating that SA might also alter the mitochondrial reserve respiratory capacity in HepaRG cells **(Figure 7D)**. After 20 mM APAP, basal and maximal respiration and respiration linked to ATP production and spare respiratory capacity were particularly reduced in HepaRG cells cultured with PA and SA **(Figure 7A-B)**. In contrast, these different respiratory parameters were about the same between HepaRG cells incubated with LA and cells treated without FAs **(Figure 7 A-B)**. Altogether, these data suggested that higher APAP-induced cytotoxicity observed with PA and SA (**Figure 5D**) could be explained at least in part by more severe mitochondrial dysfunction.

**Figure 7:**
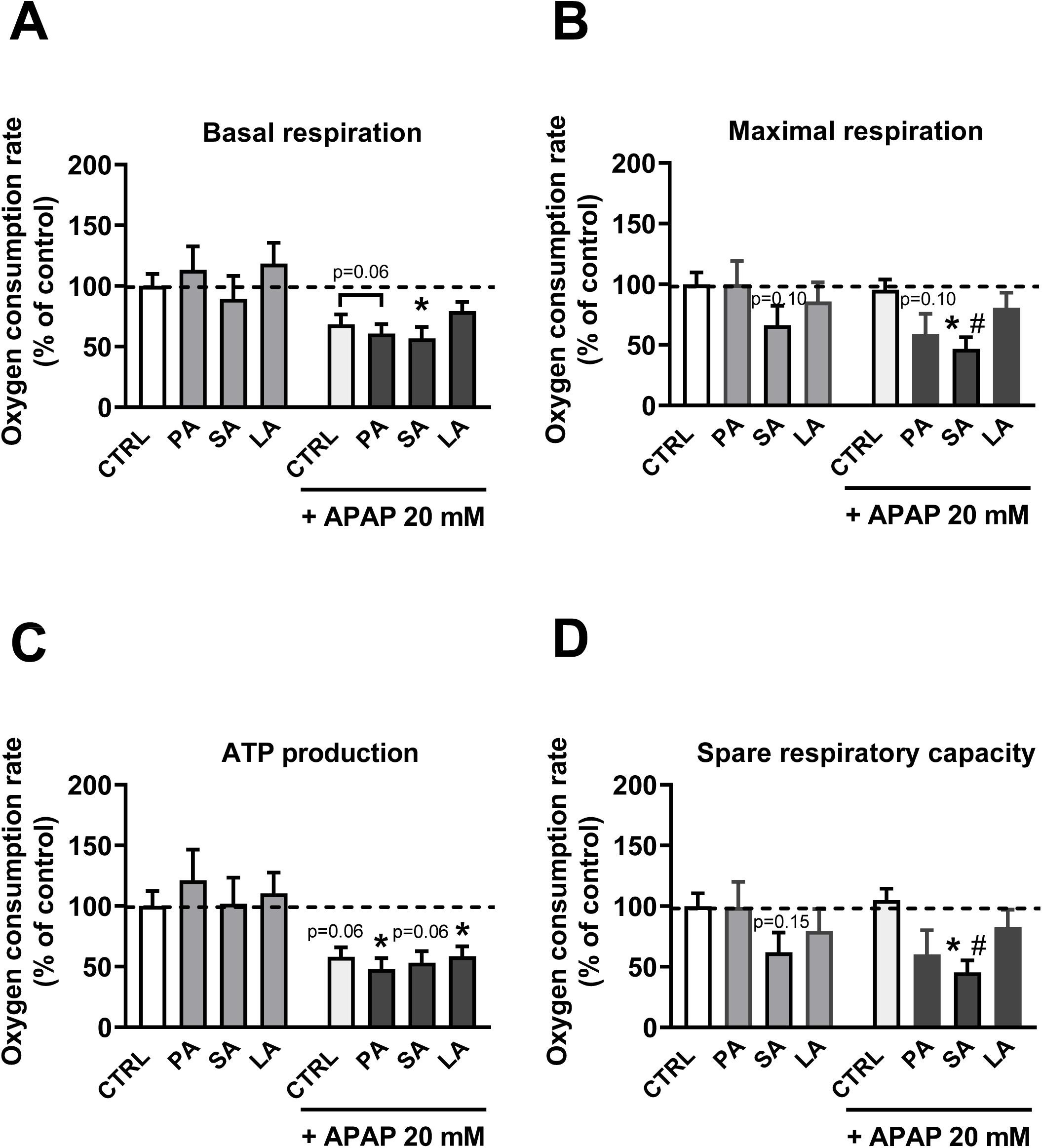
Mitochondrial respiration in HepaRG cells treated for 1 week with palmitic, stearic or linoleic acids and exposed or not to acetaminophen. Differentiated HepaRG cells were incubated for 1 week with 150 µM of palmitic (PA), stearic (SA) or linoleic (LA) acids and exposed or not to 20 mM acetaminophen (APAP) for the last 24 h of the treatment. The oxygen consumption rates (OCR) corresponding to basal and maximal mitochondrial respiration and respiration linked to ATP production as well as spare respiratory capacity were provided by the XF Cell Seahorse Mito Stress Test. Results are expressed as mean ± SEM from 5 independent cultures. * significantly different from control cells (CTRL) (p≤0.05). # significantly different from HepaRG cells treated with APAP and without fatty acids (p≤0.05).

## DISCUSSION

MASLD is the leading cause of chronic liver diseases worldwide and can evolve into more severe forms such as MASH, fibrosis and cirrhosis. Induction of hepatic CYP2E1 has been raised as an important metabolic driver in the worsening of fatty liver to MASH (Aubert et al., 2011; Abdelmegeed et al., 2012). Numerous experimental studies have proposed that hepatic CYP2E1 induction in the context of obesity and MASLD could result in part from the action of specific FAs (Aubert et al., 2012; Michaut et al., 2016; Osabe et al., 2008; Raucy et al., 2004; Sung et al., 2004). In obesity-associated MASLD, hepatic induction of CYP2E1 has been suggested as a possible adaptative mechanism to limit fat accretion in the liver by promoting ω-oxidation of FAs, thus reinforcing the β-oxidation of FAs in mitochondria and peroxisomes (Reddy et al., 2006). Besides its implication in MASLD pathophysiology, greater activity of hepatic CYP2E1 has also been implicated in the higher susceptibility to acute APAP hepatotoxicity in the setting of obesity and MASLD (Michaut et al., 2014; Begriche et al., 2023). Here, we showed that PA, SA and LA were inducers of CYP2E1 activity in differentiated HepaRG cells with only LA being steatogenic. Furthermore, PA and SA exposure aggravated APAP-induced cytotoxicity, possibly by favoring mitochondrial dysfunction.

The current study performed in HepaRG cells aimed at identifying new FAs as inducers of CYP2E1 and at determining whether these FAs could induce greater APAP cytotoxicity. Differentiated HepaRG cells were treated for 1 week with 9 FAs differing from each other by their carbon chain length and their degree of unsaturation. This broad screening led to the identification of PA, SA and LA as inducers of CYP2E1 activity in HepaRG cells. It is not surprising that only 3 out of the 9 FAs tested induced CYP2E1 activity as previous studies have highlighted that hepatic CYP2E1 induction may differ depending on the type of FAs used for cell lipid overload (Sung et al., 2004; Michaut et al., 2016). Indeed, investigations performed in differentiated HepaRG cells incubated for 1 week with 100 µM of SA or OA showed that only SA induced CYP2E1 activity in low insulin conditions whereas OA had no effect (Michaut et al., 2016). In HepG2 cells, increased CYP2E1 protein level was observed after 12 hours of LA or OA incubation whereas an exposition to ALA and DHA induced no change (Sung et al., 2004). Besides confirming our previously published data on SA-mediated CYP2E1 induction in differentiated HepaRG cells (Michaut et al., 2016), the present study identified PA and LA as two new additional FAs increasing CYP2E1 activity in these cells. Moreover, it is interesting to note that enhanced CYP2E1 activity was greater with PA and SA in comparison to LA, thus suggesting a more potent effect of long-chain saturated FAs compared to long-chain polyunsaturated FAs. This hypothesis is even more relevant considering the absence of CYP2E1 induction in response to ARA, a FA deriving directly from LA. Yet, PA, SA and LA had no effect on CYP2E1 gene expression while these 3 FAs were able to slightly increase CYP2E1 protein expression.

Since the exact mechanisms whereby FAs could induce CYP2E1 still remained unknown, we conducted a series of molecular investigations to determine possible regulatory mechanisms. Regulation of CYP2E1 expression and activity is complex and relies on several mechanisms such as transcriptional activation, increased mRNA stabilization or translation and decreased protein degradation (Koop and Tierney, 1990; Novak and Woodcroft., 2000). Our results suggested that PA, SA or LA-mediated CYP2E1 induction may result in decreased protein degradation and/or better protein stabilization rather than in increased transcription of the CYP2E1 gene. In this study, CYP2E1 regulation by FAs seems thus to rely more on post-translational mechanisms promoting CYP2E1 activity. One of the most important regulatory mechanisms for CYP2E1 is mainly *via* the stabilization of CYP2E1 protein by its own substrates which often prevents its degradation from proteasome (Song et al., 1987; Eliasson et al., 1990). Thus, further studies will be required to determine whether PA, SA and LA could impair CYP2E1 proteasomal degradation. In addition, it would be interesting to study CYP2E1 phosphorylation, which could directly modulate CYP2E1 activity (Oesch-Bartlomowicz et al., 1998).

CYP2E1 activity has been correlated with the degree of fatty liver in patients with MASLD (Emery et al., 2003; Chtioui et al., 2007). Here, we reported in HepaRG cells that only LA promoted an increase in the intracellular levels of triglycerides, while PA and SA were unable to induce steatosis. Thus, FA-mediated induction of CYP2E1 activity seemed to be independent of intracellular triglyceride accumulation. It is possible that FA-mediated increased CYP2E1 activity could be linked to the presence of certain FA derivatives able to stabilize the CYP2E1 protein, as previously mentioned, or be associated with the activation of some signaling pathways. Interestingly, hepatic CYP2E1 activity is increased in female db/db mice but not in female ob/ob mice despite more pronounced steatosis in these animals (Aubert et al., 2012). These two rodent models of obesity and type 2 diabetes present striking differences in hepatic inflammatory profile and circulating lipidome (Suriano et al., 2021), which could impact CYP2E1 activity.

As lipid homeostasis in hepatocytes is regulated by fatty acid uptake, *de novo* lipogenesis, mitochondrial β-oxidation and VLDL secretion (Ipsen et al., 2018), we investigated these four major metabolic pathways in HepaRG cells. Overall, our results suggested that LA-induced steatosis could mainly result from increased FA uptake due to higher *FAT/CD36* expression. This is in line with experimental and clinical investigations reporting that increased expression of FAT/CD36 gene and/or protein enhances FAs uptake and fat accumulation in the liver (Koonen et al., 2007; Greco et al., 2008; Miquilena-Colina et al., 2011). Furthermore, *FAT/CD36* gene expression is often increased in patients diagnosed with fatty liver or MASH in comparison to healthy individuals (Greco et al., 2008; Zhu et al., 2011). Interestingly, our targeted lipidomic data indicated that LA was highly incorporated into intracellular triglycerides following one week of treatment, suggesting that LA is preferentially used for storage. In contrast, HepaRG cells cultured with PA or SA did not induce lipid accumulation, possibly due to decreased FA uptake, increased mtFAO and reduced FA esterification into triglycerides. Interestingly, a study conducted in 3T3-L1 adipocytes exposed to various FAs showed that only SA was able to significantly reduce *FAT/CD36* gene expression indicating that regulation of the *FAT/CD36* gene may differ depending on the type of FAs (Yang et al., 2007). Conversely, *FAT/CD36* expression was significantly increased when 3T3-L1 cells were incubated with 2-bromopalmitate, a non-metabolizable FA analog (Yang et al., 2007). Thus, intracellular accumulation of unprocessed FAs could function as a signal triggering the induction of *FAT/CD36* gene expression. In our study, one hypothesis might be that increased expression of *FAT/CD36* could result from the non-accessibility of LA to oxidation and/or export since this FA is preferentially forwarded toward triglyceride storage. Reduced expression of *FAT/CD36* in HepaRG cells supplemented with PA or SA might be favored by their greater metabolizing properties as mtFAO was significantly increased in HepaRG cells treated with these two FAs.

In obesity-associated MASLD, FA oxidation could be either increased, decreased or unchanged during MASLD depending on the severity of the disease (Begriche et al., 2013; Morio et al., 2021). Activation of mtFAO was described as an adaptative mechanism to limit fat accumulation during the first stage of MASL, and as MASL progresses FAO declines gradually (Koliaki et al., 2015). As mentioned above, enhanced mtFAO oxidation in HepaRG cells cultured with PA or SA limits intracellular lipid accumulation preventing the occurrence of steatosis. Regulation of mtFAO is complex and could involve different mechanisms and factors (Houten et al., 2016). Polyunsaturated FAs have long been described as good activators of mtFAO by promoting the expression of PPARα and its target genes (Jump et al., 2005). Conversely, saturated FAs were reported as less efficient to enhance mtFAO (Jump et al., 2005; Jump., 2008). In the present study, we do not confirm these observations since PA and SA rather increase mtFAO in HepaRG cells while LA has no effect. Indeed, levels of mtFAO in HepaRG cells incubated with LA were similar to the levels measured in control cells. Moreover, no variation in expression of *PPARα* or its target genes was observed. These discrepancies could be explained by the different experimental models used in studies (*in vivo* vs *in vitro*; various cell models; different duration for FA incubation) and the availability of FAs. Regarding this latter hypothesis, *in vitro* investigations conducted on primary rat hepatocytes cultured with OA or EPA indicated an increased mtFAO in response to EPA and unchanged mtFAO in response to OA (Pawar and Jump., 2003). In this study, assessment of metabolic fate of these FAs allowed to show that incorporation of OA into neutral lipids prevents its oxidation whereas EPA barely assimilated into triglycerides were more accessible to oxidation. In our study, LA is mainly incorporated into triglycerides as previously mentioned in contrast to PA and SA. Therefore, PA- and SA-induced mtFAO could be favored by the accessibility of these two FAs to oxidation whereas unchanged mtFAO with LA could be due to a preferential use towards storage. This last effect is in line with decreased lipogenesis by LA exposure as a metabolic adaptation to excess exogenous lipid supply.

CYP2E1 induction might be involved in the greater risk of acute APAP-induced hepatoxicity in the context of obesity and MASLD (Michaut et al., 2014; Massart et al., 2022; Begriche et al., 2023). Indeed, clinical and experimental studies reported increased acute APAP toxicity in obesity and MASLD (Corcoran et al., 1987; Aubert et al., 2012; Kučera et al., 2012; Piccinin et al., 2019; Shi et al., 2019, Nguyen et al., 2008; Myers and Shaheen 2009; Chomchai and Chomchai., 2018), which are metabolic diseases associated with an induction of hepatic CYP2E1 activity (Aubert et al., 2011; Song et al., 2015; Massart et al., 2022; Begriche et al., 2023). Our experimental model of FA exposure showed that increased CYP2E1 activity was not systematically leading to higher APAP cytotoxicity. While PA and SA treatment exacerbated APAP-induced toxicity, LA was without effect compared to APAP alone. The additive effect of SA and APAP exposure on cell viability was previously reported (Michaut et al., 2016). Of note, CYP2E1 activity in HepaRG cells was increased to a lesser extent after LA exposure than PA or SA which could have had less impact on APAP metabolism and subsequent mitochondrial toxicity. In keeping with this assumption, mitochondrial dysfunction was less severe in HepaRG cells cultured with LA as compared with PA and SA. In addition, the incorporation of LA into neutral lipids might have curbed APAP-induced mitochondrial dysfunction and cell death. Previous investigations showed that esterification of unsaturated FAs into triglycerides protected cells from hepatoxicity (Listenberger, 2003). In contrast, it is well known that saturated FAs are more hepatotoxic than unsaturated species. For instance, *in vitro* investigations showed that monounsaturated fatty acids such as palmitoleate and OA are less cytotoxic than saturated FAs such as PA or SA (Wei et al., 2009; Akasawa et al., 2010; Pardo et al., 2015).

Finally, we cannot exclude that CYP2E1 subcellular localization could play a role in higher APAP-induced cytotoxicity in HepaRG cells cultured with PA and SA. Indeed, CYP2E1-mediated APAP metabolism to NAPQI and ROS overproduction can occur both within the endoplasmic reticulum and in mitochondria in accordance with its dual localization (Massart et al., 2022). Thus, CYP2E1 increased activity mediated by PA or SA might be more located at the mitochondria, which could favor mitochondrial dysfunction *via* the local production of NAPQI, which principally targets mitochondrial proteins (Ramachandran and Jaeschke., 2019; McGill and Hinson., 2020). Further investigations are needed to better characterize NAPQI production at the subcellular level as well as the exact contribution of mitochondrial CYP2E1 in the observed effects.

In summary, our study uncovered new FAs inducing CYP2E1 activity in HepaRG cells. PA- and SA-mediated CYP2E1 induction exacerbated APAP-induced mitochondrial dysfunction and cytotoxicity without inducing steatosis while LA-mediated increased CYP2E1 activity did not favor APAP toxicity despite triglyceride accumulation. Accordingly, FA-mediated increased CYP2E1 activity in HepaRG cells does not necessarily require triglyceride accumulation. Moreover, FA-mediated CYP2E1 induction does not systematically cause higher APAP-induced cytotoxicity. By promoting steatosis, LA might curb APAP-induced mitochondrial dysfunction and cell death. We believe that our investigations provide new knowledge about the regulation of CYP2E1 by some FAs, which might help to find nutritional or pharmacological strategies aimed at limiting APAP-induced toxicity in the setting of obesity and MASLD. This work also provides important information about the nature of FAs that promote CYP2E1 induction and possibly a faster progression of fatty liver to NASH.

## Acknowledgment

We are grateful to INSERM (Institut National de la Recherche et de la Santé Médicale) for its constant financial support. Clémence Penhoat was a recipient of a joint fellowship from the Region Bretagne (ARED) and INSERM. We are also grateful to master students who participated to this work (Matthieu Franc, Jules Vaxelaire and Cassandra Rayer).

---

**Supplemental figure 1.**
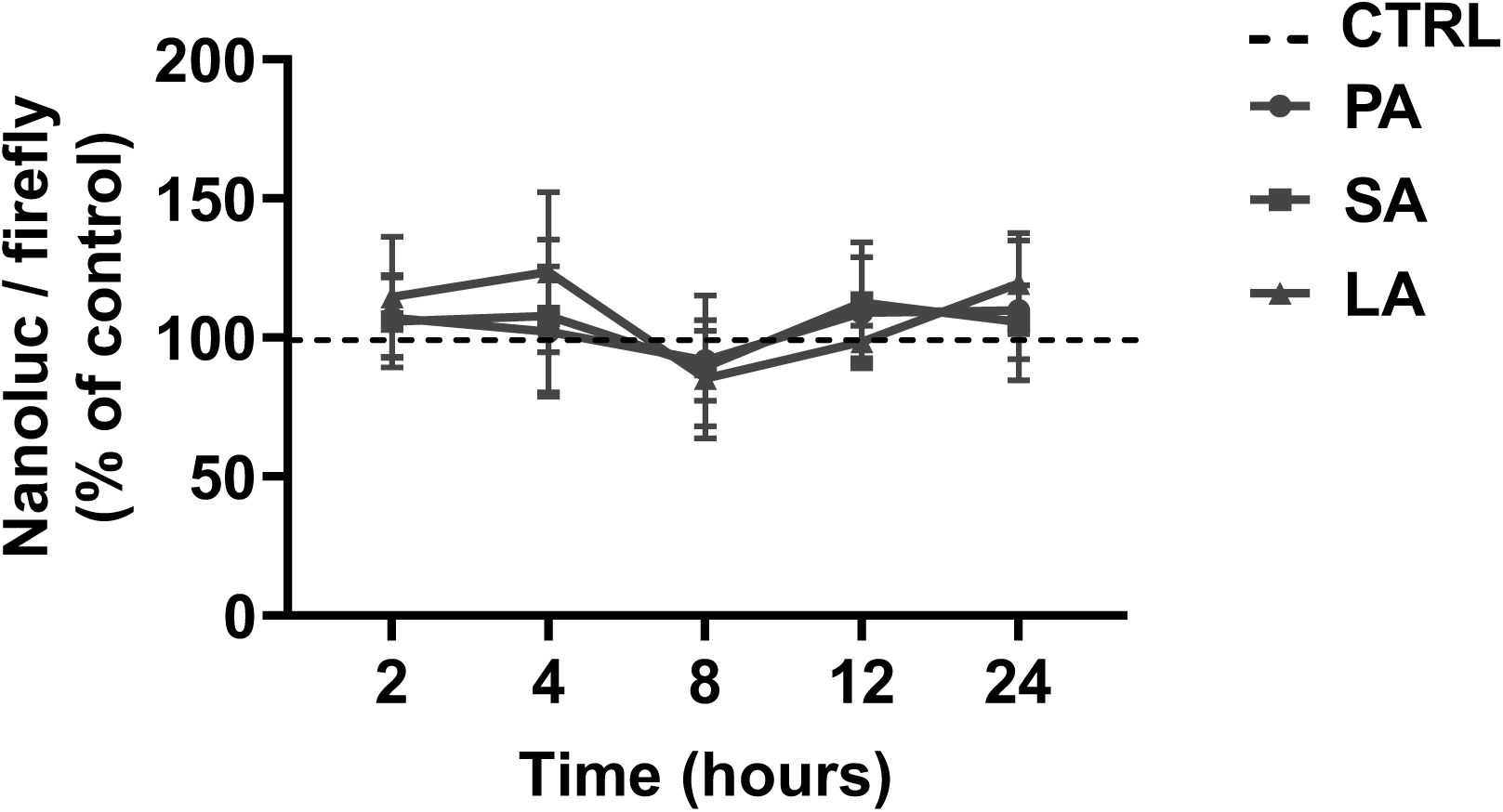

**Supplemental figure 2.**
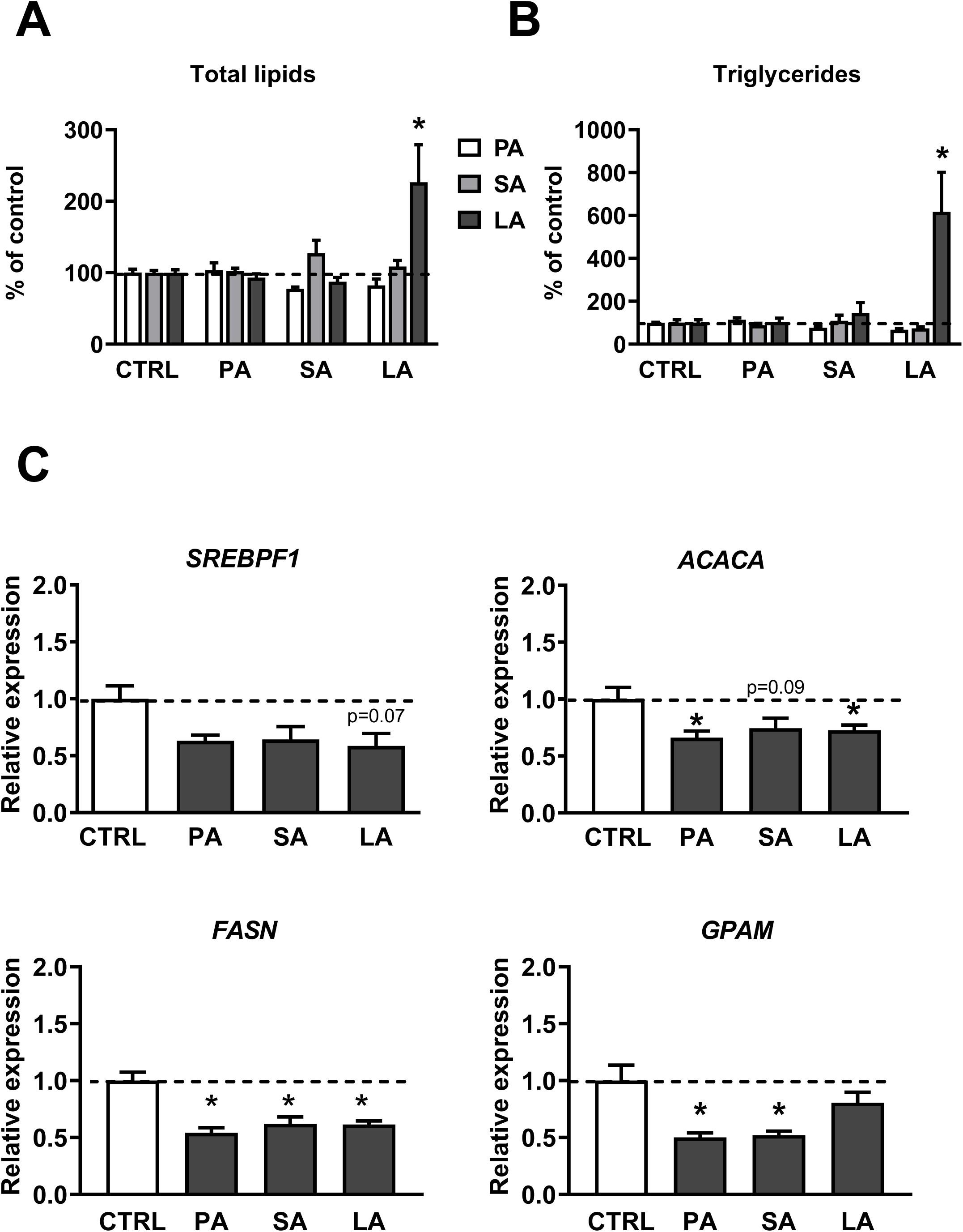

**Supplemental figure 3.**
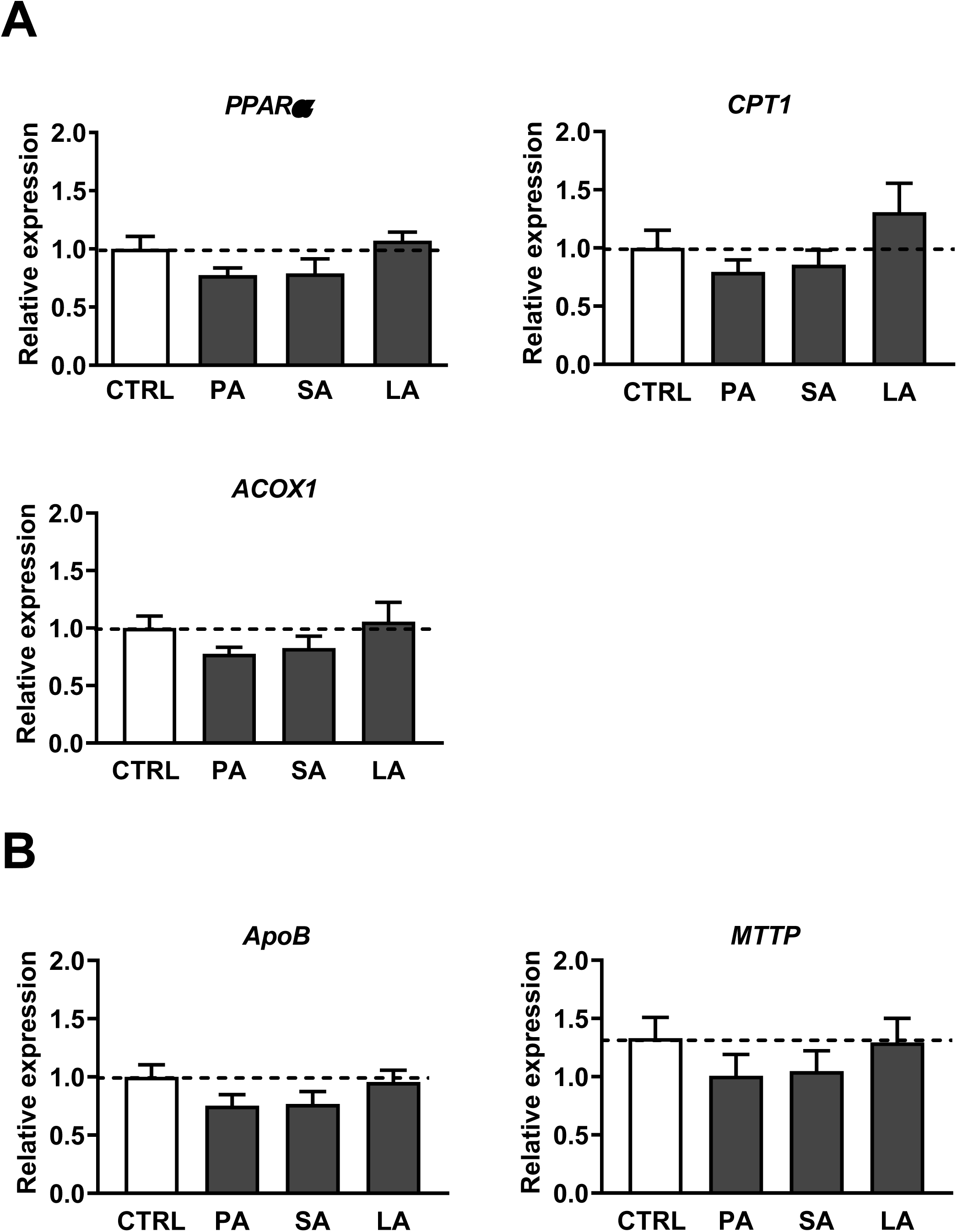

