## Supplemental data for "Fatty acid-mediated induction of CYP2E1 activity in HepaRG cells is not systematically associated with exacerbated acetaminophen cytotoxicity"

### Supplementary data

#### Supplementary Materials and Methods

##### *Fatty acid composition of lipid and triglyceride fractions*

Hexane–isopropanol (3/2, v/v) was used to extract total lipids from HepaRG cells treated with 150 µM of palmitic (PA), stearic (SA) and linoleic (LA) acids for 1 week (Rioux et al., 2000). Lipid classes were then separated by thin layer chromatography. Glass plates (20×20 cm) were coated with SilicaGel 60H (Merck, Darmstadt, Germany) according to 0.5-mm thickness. Glass plates with samples were developed with a mixture of hexane/diethyl ether/acetic acid (85/15/1, v/v/v). Spots of interest were revealed with primuline (0.02% in acetone w/v) under UV light and then collected. To determine fatty acid composition of total lipid and triglyceride fractions, fatty acids of each fraction were transformed to Fatty Acid Methyl Esters (FAMEs). First**,** fatty acids were methylated according to a two-step procedure using NaOH (0.5 M in methanol) and $BF_3$ (12% in methanol) (Metcalfe et al., 1966) and then extracted twice with pentane and NaCl (0.9%). Then, FAMEs were analyzed with an Agilent 6890N gas chromatograph (Agilent Technologies, Santa Clara, CA, USA) equipped with a bonded fused silica capillary column (BPX70; 60 m×0.25 mm, 0.25 µm thickness; SGE Analytical Science, Melbourne, Australia). The temperature program started at 150°C, then increased to 250°C at 4°C/min and finally held for 2 min. Helium was used as carrier gas. Mass spectra were recorded with an Agilent 5975C (Agilent Technologies, Santa Clara, CA, USA) operated under electron impact ionization conditions (electron energy 70 eV, source temperature 230°C). Mass spectrum of peaks was carefully checked to ensure proper identification. Peak integration was accomplished with MassHunter WorkStation Software Qualitative Analysis vB.07.00 (Agilent Technology, Santa Clara, CA, USA). Fatty acid data were handled as a percentage of total fatty acids and then expressed as percentage of control.

***Cloning of CYP2E1 promoter and transcriptional assay***

Genomic DNA fragment corresponding to HepaRG CYP2E1 promoter gene sequence was amplified by PCR using 2x Platinum™ SuperFi™ PCR Master Mix kit (Thermofisher Scientific) with the forward primer 5' TGGCCTCGGCGGCCAAGCTTCCCTCGGGGCCTCAGGGAGCCGCAGCC 3' and the reverse primer 5'TCTTCGAGTGTGAAGACCATGGTGCCGCTGGGGCCCTGCTGCCAG 3'. PCR products were purified on agarose gel and cloned into the pNLucP reporter vector (Promega) using the NEBuilder HiFi DNA Assembly Master Mix kit (NEB). A sequencing was performed (Eurofins) in order to ensure the proper amplification and insertion of CYP2E1 promoter gene into the pNLucP reporter vector. Prior to transcriptional assay, HepG2 cells were plated 48 h before transfection in 24-well culture plates at $5 \times 10^4$ cells/cm$^2$ and incubated in DMEM medium supplemented with 10% FBS, 100 units/mL penicillin, 100 µg/mL streptomycin, 5 µg/mL insulin, and 50 µM hydrocortisone hemisuccinate. Lipofection was performed using the pCYP2E1NLucP reporter vector. RedFLuc plasmid was used as a normalization vector. Lipofection complexes were prepared in OptiMEM Reduced Serum Medium with 200 ng of pCYP2E1nLuc, 2 ng of RedFLuc (pCYP2E1nLuc:RedFLuc ratio of 100:1) and 4 µL of ViaFect™ (plasmids:ViaFect™ ratio of 1:2) per well, kept at room temperature for 20 minutes, and added dropwise on the cells. Media was renewed the day after transfection to stop the reaction. HepG2 cells were subsequently treated with palmitic (PA), stearic (SA) or linoleic (LA) FAs for 2h, 4h, 8h, 12h or 24h. Luciferase measurement was then performed using the NanoGlo Dual Luciferase Reporter Assay System kit (Promega). CYP2E1 promoter activity was expressed as luminescent signal of Nanoluc/ Firefly. This model was validated using IL4 dose-response, given the known inductive effects of this cytokine on CYP2E1 promoter as previously described (Abdel-Razzak et al., 2004). HepG2 cells were used at passage 13 to 30.

### Supplementary figure legends

**Supplementary Figure 1: CYP2E1 promoter activity in HepG2 cells treated with palmitic, stearic and linoleic acids**

HepG2 cells transfected with pCYP2E1NLucP reporting vector were treated with 150 µM palmitic (PA), stearic (SA) and linoleic (LA) FAs for 2, 4, 8, 12 and 24h. CYP2E1 promoter activity was determined by luciferase assay. Results are expressed as mean ± SEM from 4 independent cell cultures. The horizontal dashed line represents the level of control cells (CTRL).

**Supplementary Figure 2: Fatty acid composition of intracellular lipids and lipogenic gene expression in HepaRG cells treated for 1 week with palmitic, stearic or linoleic acids.**

Differentiated HepaRG cells were incubated for 1 week with 150 µM palmitic (PA), stearic (SA) or linoleic (LA) FAs.  Fatty acid composition of total lipid (A) and triglyceride (B) fractions determined by GC-MS (n=4). (C) Expression of different genes involved in *de novo* lipogenesis (n=7). Results are expressed as mean ± SEM. n indicates the number of independent cell cultures. The dashed line represents the level of control cells (CTRL). * significantly different from control cells (CTRL) ($p \leq 0.05$).

**Supplementary Figure 3: Expression of different genes involved in fatty acid oxidation and VLDL secretion in HepaRG cells treated for 1 week with palmitic, stearic or linoleic acids.**

Differentiated HepaRG cells were incubated for 1 week with 150 µM of palmitic (PA), stearic (SA) or linoleic (LA) FAs.  Expression of different genes involved in fatty acid oxidation (A) and VLDL secretion (B) (n=7). Results are expressed as mean ± SEM. n indicates the number of independent cell cultures. The dashed line represents the level of control cells (CTRL).

#### Table 1

| Gene symbol (and alias) | Gene name | Accession number | Forward primer (5'-3') | Reverse primer (5'-3') |
|---|---|---|---|---|
| *ACACA (ACC1)* | Acetyl-CoA carboxylase alpha | NM_198839.2 | GGTGGATCGGAGATTTCATAGAG | AGGCTCCAGATGACGATAGA |
| *ACOX1* | Acyl-CoA Oxidase 1 | NM_001185039.2 | AGCCAGCGTTATGAGGTG | ATGCCCAAGTGAAGATCCAG |
| *APOB* | Apolipoprotein B | NM_000384.3 | AAGTATGGGATGGTAGCACAAG | TGGAGGTGATGTGGATTTGG |
| *CYP2E1* | Cytochrome P450 family 2 subfamily E member 1 | NM_000773.4 | GAACTTCCACCTACTCAGCAC | CTCCTTCACCCTTTCAGACAC |
| *CPT1A (L-CPT1)* | Carnitine palmitoyltransferase 1A | NM_001876.4 | CGGGAGGAAATCAAACCAATTC | CTGGGATCCGGGAAGTATTAAA |
| *CD36 (FAT)* | CD36 molecule | NM_001001548.3 | GCCAGGTATTGCAGTTCTTTTC | TGTCTGGGTTTTCAACTGGAG |
| *Cyclophilin B* | CycloB | NM_000942.5 | CAGTGGATAATTTTGTGGCCTTAG | CTCACCGTAGATGCTCTTTCC |
| *FASN* | Fatty acid synthase | NM_004104.5 | CTGAGCAGTACACACCCAAG3 | GATACTTTCCCGTCGCATACC |
| *GPAM (GPAT1)* | Glycerol-3-phosphate acyltransferase, mitochondrial | NM_020918.6 | CTGTGCTACCTTCTCTCCAAT | ATATCTTCCTGGTCATCGTGC |
| *MTTP* | Microsomal triglyceride transfer protein | NM_000253.3 | TACCAGGCTCATCAAGACAAAG | CTGACACCCAAGACCTGATTT |
| *PLIN1* | Perilipin 1 | NM_002666.5 | GCCATGTCCCTATCAGATGC | GTTGTCGATGTCCCGGAATT |
| *PLIN2* | Perilipin 2 | NM_001122.4 | GCTCCATTCTACTGTTCACCTG | CTCCTTTTCCACTCTACCCATG |
| *PPARα* | Peroxisome proliferator activated receptor alpha | NM_001001929.3 | GTTCTGGAAGCTTTGGCTTTAA | GAAAGCGTGTCCGTGATGA |
| *SREBP1F (SREBP1c)* | sterol regulatory element binding protein 1c | NM_001321096 | GAGCCATGGATTGC | GGAAGTCACTGTCTTGGTTGTTGA |
